# Weight cycling-induced hypothalamic and metabolic tissue immune remodeling is uncoupled from metabolic dysfunctions

**DOI:** 10.1101/2025.06.27.661006

**Authors:** Hendrik J.P. van der Zande, Alon Zemer, Frank Vrieling, Lisa Smeehuijzen, Brecht Attema, Habib Muallem, Jenny Jansen, Anouk van der Linden, Sivan Prass, Daniëlle Wessels, Meirong Ren, Yulia Haim, Evert M. van Schothorst, Sander Kersten, Assaf Rudich, Rinke Stienstra

**Author notes:** Corresponding author: Rinke Stienstra, Stippeneng 4, 6708WE Wageningen, **+**31317482338.

## Abstract

**Background:** Obesity-induced insulin resistance is associated with white adipose tissue (WAT) and liver inflammation, which are both mitigated by weight loss. However, most individuals undergoing weight loss will regain lost weight, resulting in weight cycling (WC), which may exacerbate metabolic dysfunctions. Here, we studied the immunometabolic impact of WC in mice.

**Methods:** C57BL/6J mice were exposed to two cycles of weight gain and weight loss by alternating between low (LFD) and high-fat diet (HFD) feeding. Animals were sacrificed when WC mice were weight stable for 10 weeks upon weight loss (WC-lean) and after a subsequent exposure to weight regain for 10 weeks (WC-obese), and compared to mice persistently fed LFD (LFD-lean) or HFD (HFD-obese).

**Results:** Body weight stabilized at a higher level in WC-lean mice after two weight gain/loss cycles compared to LFD-lean controls. While insulin resistance, metabolic tissue inflammation, and hepatic steatosis normalized between the two groups of lean mice, WC-lean mice exhibited features of WAT dysfunction. In the hypothalamus, inflammatory microglia were less abundant in WC-lean mice compared to LFD-lean mice, but mean individual microglial cell volume was larger. WC-obese mice stabilized at slightly lower weight compared with HFD-obese controls. Intriguingly, WC-obese mice exhibited increased WAT macrophages and reduced WAT and liver effector T cells compared to HFD-obese mice, whereas energy intake, body composition, whole-body insulin resistance and hepatic steatosis were similar.

**Conclusions:** Our results suggest that WC in mice differently impacts animals in the weight-stable lean and obese states. WC-lean mice display features of a novel body weight settling point, associated with hypothalamic inflammatory changes. However, metabolic dysfunctions were uncoupled from WC-induced metabolic tissue inflammation in WC-obese mice.

## Introduction

A chronic imbalance favoring energy intake over expenditure results in overweight and obesity, which predisposes to sequelae such as type 2 diabetes, metabolic dysfunction-associated steatotic liver disease (MASLD), cardiovascular diseases, neurodegenerative diseases, and certain types of cancer (1). Body weight loss by any intervention is an effective strategy for reducing the risk and burden of these diseases (2–6). However, maintaining a lower body weight remains a considerable challenge for most, both after intensive lifestyle interventions (7–9) and discontinuation of obesity pharmacotherapy (10–12). This phenomenon of weight cycling (WC), or yoyo-dieting, has been reported to abrogate the metabolic benefits of previous weight loss and may even aggravate obesity-associated dysfunctions (12, 13). The growing number of individuals at risk for WC due to the expanding toolkit of obesity treatments, and the potential of WC to impair metabolic health, underscore the urgency to unravel its mechanisms.

Obesity profoundly changes the immune cell composition of white adipose tissue (WAT) and liver, which is linked to metabolic tissue inflammation and the development of insulin resistance (14). Indeed, lean, insulin-sensitive WAT is characterized by a high abundance of eosinophils and type 2 innate lymphoid cells (ILC2s) (15, 16). By contrast, obese adipose tissue is characterized by increased monocyte chemoattractant protein 1 (MCP1)-driven monocyte recruitment and their development into pro-inflammatory lipid-associated macrophages (LAMs), driving an inflammatory milieu that associates with impaired insulin sensitivity (17–19). Similarly, in the liver, obesity results in the loss of resident Kupffer cells (KCs), which are replaced by pro-inflammatory monocyte-derived Kupffer cells (moKCs). These moKCs have been suggested to contribute to MASLD (20, 21).

The concept of immunological memory has been classically associated with responses to infectious pathogens and vaccines. Recently, it was shown to also encompass non-infectious stimuli. Indeed, it is suggested that metabolic events, such as hyperglycemia, can induce trained immunity, thereby influencing secondary inflammatory and metabolic responses (22). Interestingly, several studies indicate that obesity-induced inflammatory characteristics are sustained after weight loss, despite improvements in metabolic homeostasis (23–25), supporting the existence of immunological obesogenic memory. Given the importance of the brain in regulating energy intake and expenditure (26, 27), such obesogenic memory may involve both peripheral and central (brain) adaptations. However, important gaps in our understanding remain: (i). How does WC-induced obesogenic memory manifest in the lean and obese states? (ii). Will obesogenic memory fade with prolonged weight stabilization? (iii). Does a history of WC exacerbate inflammation and metabolic dysfunction upon subsequent weight regain?

To address these outstanding questions, we established a murine model of *ad libitum* diet-induced WC. We assessed obesogenic memory by comparing lean WC mice (WC-lean) after extended weight loss and weight stabilization to persistently low-fat diet (LFD)-fed lean controls (LFD-lean), and obese WC mice (WC-obese) after an extended phase of weight regain and weight stabilization to persistently high-fat diet (HFD)-fed obese controls (HFD-obese).

## Materials & Methods

### Animals and model

All experiments followed the Guide for the Care and Use of Laboratory Animals of the Institute for Laboratory Animal Research and were approved by the Central Authority for Scientific Procedures on Animals (CCD, AVD10400202115283) and the Institutional Animal Care and Use Committee of Wageningen University (2021.W-0016.002). Six-week-old wild-type C57BL/6JOlaHsd male mice (n=90) were purchased from Envigo (the Netherlands) and were acclimatized to the animal facility for 4 weeks before the start of the experiment. Animals were singly housed due to fighting among cage-mates, in a temperature-controlled (21°C) room with a 12-hour light-dark cycle, with *ad libitum* access to food (synthetic low-fat diet [LFD]; 10% kCal derived from lard fat, D12450H, Research Diets), tap water and cage enrichment in the form of a tunnel and nesting material (tissues).

The weight cycling model was adapted from (28). This included an LFD-fed persistently-lean control group (LFD-lean), a high-fat diet (HFD, 45% kCal derived from lard fat, D12451, Research Diets)-fed persistently-obese control group (HFD-obese), and a weight-cycled (WC) group that underwent three cycles of obesity as follows (**Figure 1A**):

1. Initial weight gain via HFD feeding for four weeks;
2. Weight loss via LFD feeding, until body weight difference between the weight cycling group and the lean control group was statistically insignificant;
3. Weight regain via HFD feeding, until body weight difference between the weight cycling group and the obese control group was statistically insignificant;
4. Weight loss via LFD feeding, until WC mice were weight stable for 10 weeks;
5. Final weight regain via HFD feeding for 10 weeks.

**Figure 1:**
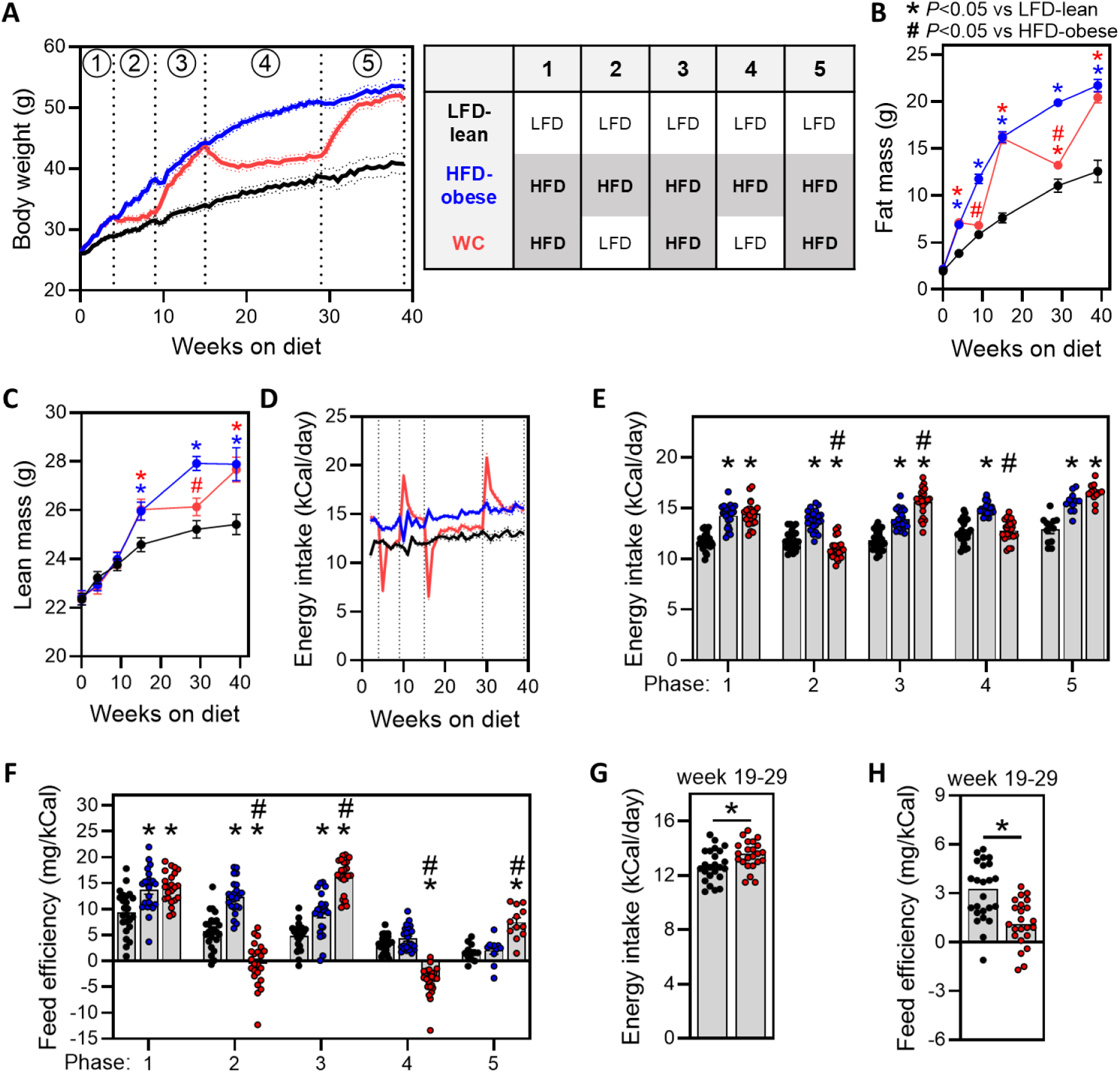
Weight cycling affects body weight regulation. (**A**) Left panel: Body weight development of the LFD-lean control group (black), HFD-obese control group (blue) and weight cycled (WC) group (red). Right panel: Overview of which diet was fed per group in each phase of the study. (**B-C**) Fat mass (B) and lean mass (C) measured at every diet switch. (**D-E**) Energy intake over time (D) and per phase (E) of the experiment. (**F**) Body weight gain and energy intake were used to calculate feed efficiency, shown per phase of the experiment. (**G-H**) Average energy intake (G) and feed efficiency (H) were determined for the LFD-lean and WC groups during the weight-stable phase (week 19-29). Data are expressed as mean ± SEM (n=23-24 mice per group until week 29, and n=11-12 mice per group from week 29-39). Significance was tested by Two-Way ANOVA with Tukey correction for multiple testing (B, C, E, F) or independent samples t-test (G, H). * *p* < 0.05 vs LFD-lean, # *p* < 0.05 vs HFD-obese.

Before every new group allocation, groups were standardized based on body weight, fat mass and fasting blood glucose to ensure no *a priori* differences that may confound the data. Since the body weight response of C57BL/6J mice to HFD feeding is markedly variable, and since body weight was an important variable in the study design, we continued with the mice within the 10-90^th^ percentile of body weight after 4 weeks on either LFD or HFD. Half of the mice were sacrificed after two full weight cycles when WC mice were weight stable for 10 weeks, before the last 10-week re-exposure to HFD (n=11-12 mice per group). Sample size was determined via an *a priori* power calculation, allowing to detect a relevant difference in HOMA-IR of 15%. Mice were sacrificed through cervical dislocation following isoflurane anesthesia. Researchers were not blinded to the experimental groups on test days, yet were blinded during subsequent analyses.

### Food intake and body composition

Body weight and food intake were determined every week throughout the experiment by weighing the mice and food pellets. Feed efficiency was calculated from body weight gain and energy intake and expressed as mg body weight per kCal food. Body composition was determined in conscious, unrestrained mice using an EchoMRI (Echo Medical Systems) at every diet switch (indicated with the dashed vertical lines in **Figure 1A**). At sacrifice, visceral white adipose tissue (epididymal; eWAT), subcutaneous white adipose tissue (inguinal; iWAT), and liver were weighed and collected for downstream processing.

### Plasma analysis

Fasted blood samples were collected one week before sacrifice from the tail vein of 5-hour-fasted mice using glass capillaries, and fasting blood glucose was determined using a hand-held glucometer (Glucofix Tech; A. Menarini). Plasma insulin (Chrystal Chem #90080), alanine transaminase (ALT) activity (Sigma-Aldrich #MAK052), triglycerides (Instruchemie #10720P), total cholesterol (Sopachem #1 1350 99 10 021) and non-esterified fatty acids (Instruchemie #3055) were determined using commercially available kits per manufacturer’s instructions. The homeostatic model assessment of insulin resistance (HOMA-IR) adjusted for mice (29) was calculated as follows: ([glucose (mg/dl)*0.055]*[insulin (ng/ml)*172.1])/3857.

### Liver triglyceride content

∼15 mg liver samples were homogenized in 10 mM Tris-base containing 250 mM sucrose and 2 mM EDTA (pH 7.5) at 2% w/v using a Tissue Lyzer. Liver triglyceride content was subsequently determined using a commercially available kit (Instruchemie #10720P) according to the manufacturer’s instructions.

### RNA-isolation and RT-qPCR

RNA was isolated from snap-frozen eWAT, eWAT adipocytes and liver samples using TRIzol RNA-Isolation reagent (Invitrogen) and the phenol/chloroform RNA extraction method. Total RNA (0.5-1 μg) was reverse transcribed using the iScript cDNA synthesis kit (Bio-Rad). Real-time qPCR was performed on a CFX384 Touch Real-Time PCR detection system (Bio-Rad) using the SensiMix SYBR No-ROX kit (Bioline). Gene expression was determined by calculating starting quantities using the standard curve method, normalizing starting quantities to the housekeeping gene *36b4* and expressed as fold change compared to the control condition. Primer sequences are listed in **Supplementary Table 1**.

### Isolation of cells from blood, adipose tissue, and liver for flow cytometry

At sacrifice, blood was collected from the retro-orbital plexus of anesthetized mice in EDTA-coated micro sample tubes (Sarstedt) via orbital removal. 200 μL whole blood was next mixed with 280 μL Proteomic Stabilizer PROT1 (Smart tube inc.) and incubated and cryopreserved at −80°C according to the supplier’s protocol. The remaining blood was centrifuged in an Eppendorf centrifuge for 15 minutes at 5,000 rpm at 4°C and plasma was stored at −80°C.

On analysis days, cryopreserved whole blood samples were thawed at 4°C for 30 minutes, and erythrocytes lysed using 1X Thaw-Lyse buffer (Smart tube inc.) according to manufacturer’s instructions. During erythrocyte lysis, cells were filtered through 100 μm and 40 μm cell strainers (Pluriselect).

eWAT and liver samples were processed as previously described (30). Briefly, eWAT and livers of two mice from the same group were pooled and were cut into small pieces using razors. eWAT fat pads were digested in 5 mL digestion buffer containing 1 mg/mL collagenase type II from *Clostridium histolyticum* (Sigma-Aldrich) in Krebs buffer supplemented with 100 mM HEPES, 20 mg/mL BSA and 6 mM D-Glucose for 45 minutes at 37°C/5%CO_2_ while shaking (100 RPM). Digestion was stopped by adding 5 mL wash buffer (PBS supplemented with 1% FCS and 2.5 mM EDTA), and the solution was filtered through 250 μm Nitex filters (Sefar). Infranatant containing the stromal vascular fraction (SVF) was collected, and erythrocytes were lysed using ice-cold erythrocyte lysis buffer (in-house, 0.15 M NH_4_Cl, 1 mM KHCO_3_, 0.1 mM Na_2_EDTA). The remaining adipocytes were collected, frozen on dry ice and stored at −80°C. SVF cells were finally filtered through 40 μm cell strainers and washed with wash buffer (400 RCF, 10 minutes, 4°C). The liver was digested in 5 mL RPMI supplemented with 1 mg/mL collagenase V, 1 mg/mL dispase II, 1 mg/mL collagenase D, and 30 U/mL DNase I for 25 minutes at 37°C/5%CO_2_ while shaking (100 RPM). Digest was filtered through 100 μm cell strainers and washed twice with 40 mL wash buffer (300 RCF, 5 minutes, 4°C). Following erythrocyte lysis, cells were finally filtered through 40 μm cell strainers. All isolated and filtered cells were counted using a hemocytometer.

### Flow cytometry

Up to 1×10^6^ blood cells were plated in a 96-well V-bottom plate for immunophenotyping by flow cytometry. Blood samples were barcoded using anti-mouse CD45-Pacific Blue (1:200) and anti-mouse CD45-APC-Fire750 (1:800) in FACS buffer (1%BSA and 2 mM EDTA in PBS) supplemented with True-Stain Monocyte Blocker (Biolegend; to avoid non-specific antibody binding of PE- and APC-based tandem dyes by myeloid cells) and TruStain FcX (Biolegend; anti-mouse CD16/32) for 15 minutes at RT protected from light. After washing away unbound barcoding antibodies using FACS buffer, barcoded samples from two mice of the same group were pooled and processed as one. Samples were incubated for 20 minutes at RT protected from light with antibodies directed against B220, CD3, CD4, CD8a, CD11b, CD44, Ly6C, Ly6G, NK1.1, Siglec-F and TCRγδ in FACS buffer supplemented with True-Stain Monocyte Blocker and Brilliant Stain Buffer Plus (BD Biosciences; to prevent staining artifacts caused by interactions between Brilliant Violet dyes). Antibody details are listed in **Supplementary Table 2**.

0.3×10^6^ eWAT SVF cells and up to 1×10^6^ liver cells were plated in a 96-well V-bottom plate, washed with PBS and incubated with the viability dye Zombie Aqua (Biolegend) in PBS supplemented with TruStain FcX for 20 minutes at RT protected from light. Three samples were barcoded using combinations of anti-mouse CD45-Pacific Blue and anti-mouse CD45-APC-Fire750 in FACS buffer for 15 minutes at RT protected from light. Following washing and pooling samples in FACS buffer, cells were incubated with antibodies directed against B220, CD3, CD9, CD11b, CD11c, CD64, F4/80, Ly6C, MHCII, NK1.1 and Siglec-F for analysis of myeloid cells, and B220, CD3, CD4, CD8a, CD25, CD44, CD62L, NK1.1 and TCRγδ for analysis of lymphocytes in FACS buffer supplemented with True-Stain Monocyte Blocker and Brilliant Stain Buffer Plus for 15 minutes at RT protected from light. Liver cells were incubated with the same antibodies except anti-F4/80, and including anti-CLEC2 and anti-TIM4 for analysis of Kupffer cells. Cells were next fixed in either 2% formaldehyde in PBS (prepared from 16% Methanol-free formaldehyde solution, Pierce) for 15 minutes at RT protected from light (for myeloid cells) or the True-Nuclear Transcription Factor buffer set (Biolegend; for lymphocytes) according to manufacturer’s instructions. Following washing with FACS buffer, eWAT SVF samples were next incubated with HSC LipidTOX Green (Invitrogen) to stain neutral lipids for 30 minutes at RT protected from light. For analysis of myeloid cells, samples were acquired immediately, and samples for assessing lymphocyte subsets were stored protected from light at 4°C. The next day, cells were washed and permeabilized using the TrueNuclear Transcription Factor buffer set, and incubated with anti-FOXP3 in True-Nuclear permeabilization buffer for 15 minutes at RT protected from light. Following washing with permeabilization buffer and FACS buffer, samples were acquired on a CytoFLEX flow cytometer (Beckman Coulter).

### Olink plasma proteomics

Circulating inflammatory proteins were measured in plasma samples via Proximity Extension Assay technology using the Olink Target 48 panel (Olink), as per the manufacturer’s instructions.

### Hypothalamic immunostaining and microglial phenotyping

Brains were collected immediately after mice were killed, incubated overnight in 4% paraformaldehyde (PFA) and then transferred to a 30% sucrose solution. Brains were frozen in OCT blocks and sectioned into 45 μm sections using a Leica cryostat. Sections were permeabilized with 0.5% Tween in PBS (PBST), blocked with 10% donkey serum, and subsequently incubated overnight at 4°C with primary antibodies for Iba1 (1:500, goat-anti-mouse, Abcam #ab5076) and p-NFκB (1:500, rabbit-anti-mouse, Abcam #ab86299). Sections were next washed and incubated with the appropriate secondary antibodies (Iba1: donkey-anti-goat-Alexa Fluor 647, Jackson ImmunoResearch Laboratories #705-605-147; p-NFκB: donkey-anti-rabbit-Alexa Fluor 488, Abcam #ab150073; 1:250 for both). For nuclear staining, sections were stained with DAPI (1:3000) for 10 minutes before mounting. Images were obtained using a FluoView 1000 confocal microscope (Olympus). Each image contained 34-41 steps z-stacks, with 0.75 µm step size. Imaris software (Oxford instruments) 34-based stacks surfaces analysis was used for cell counting, as well as to assess cell volume, colocalization of microglial pNFkB-positive cells, and to detect nuclei.

### Oral glucose tolerance test

Whole-body glucose tolerance was determined via an oral glucose tolerance test (oGTT) at 1 week before sacrifice. In short, mice were fasted for 5 hours, after which fasting blood glucose was determined (t=0) using a hand-held glucometer and a fasted blood sample was collected in glass capillaries. 1.5 g/kg body weight of D-Glucose in PBS was next administered via oral gavage, and blood glucose was measured at 20, 40, 60, 90, 120 and 150 minutes post gavage. At 20 minutes post-gavage, an additional blood sample was collected in glass capillaries to determine glucose-induced insulin secretion.

### Histology

Pieces of liver and eWAT were fixed in 4% formaldehyde immediately upon collection. Tissues were embedded in paraffin, sectioned at 4 μm and stained with Hematoxilin and Eosin (H&E) using standard protocols. Three to five fields at 10x magnification were used for the analysis of adipocyte diameter and hepatic steatosis.

### *In vitro* 3T3-L1 nutrient cycling model

Murine 3T3-L1 cells (ATCC CL-173) were cultured and differentiated following the manufacturer’s protocol (ATCC). Once fully differentiated, 3T3-L1 adipocytes underwent nutrient cycling by culturing cells in high (25 mM) or low (1 mM) D-glucose-containing medium (DMEM supplemented with 1 mM sodium pyruvate (both Gibco), 10% FCS and 1% P/S), by alternating this medium every 3 days for 12 days. The medium of control adipocytes was refreshed on the same days as the nutrient-cycling adipocytes and were cultured in medium with 25 mM glucose for all 12 days.

Glucose concentrations in supernatants were measured using the Glucose GOD Fluid Stable kit following the manufacturer’s protocol (Diagnostic Systems). Lactate concentrations in culture supernatants were quantified through the conversion of lactate by lactate oxidase (Sigma-Aldrich #L9795) and the subsequent oxidation of Amplex Red (Invitrogen #A12222) by HRP (Thermo Fisher #31490). Prior to the reaction, proteins were first precipitated using 13.7% perchloric acid to avoid interference by lactate dehydrogenase. Lactate concentrations were determined using a lactate standard (Merck #L7022). MCP-1 concentrations in supernatants were measured using the MCP-1/CCL2 mouse ELISA kit following manufacturer’s protocol (Invitrogen #88-7391-22).

### Data analysis

Data are presented as mean ± standard error of the mean (SEM). Flow cytometry data was analyzed using FlowJo version 10.10 (BD Life Sciences). Principal component analysis (PCA) was performed in R version 4.3.2 using the mixOmics package version 6.24.0 (31). Centered log ratio transformation was applied to immune cell composition data prior to PCA, to resolve the constant-sum constraint of inherently-correlated composition data. Statistical significance was tested using GraphPad Prism 10.0 (GraphPad Software) using one-way ANOVA with Tukey correction for multiple testing or as indicated. Differences between groups were considered statistically significant at *p* < 0.05. Correlogram was generated using the corrplot package version 0.92 (32). The remaining data were visualized using GraphPad Prism 10.0.

## Results

To study the immunometabolic consequences of weight cycling, we employed a previously reported model of repetitive exposure to high-fat diet (HFD)-induced obesity (28). Mice were either persistently fed a low-fat diet (LFD; LFD-lean), HFD (HFD-obese) or exposed to weight cycling (WC) via HFD feeding alternated by LFD feeding until the weight of the respective control group, or weight stabilization, was reached (**Figure 1A**). As expected, HFD feeding resulted in increased body weight, fat mass, lean mass, energy intake and feed efficiency (body weight increase per kCal food intake) (**Figure 1A-F**). WC mice switched between LFD-lean and HFD-obese control body weight, fat mass, lean mass and energy intake (**Figure 1A-D**), as per design. Energy intake significantly changed acutely following diet switches (**Figure 1D**). Moreover, average energy intake was significantly increased and decreased in WC mice compared to the LFD-lean and HFD-obese groups, respectively, yet only during the first weight loss and weight gain phases (**Figure 1E**). Feed efficiency was modulated by WC in all weight loss and weight gain phases (**Figure 1F**).

Strikingly, following two cycles of weight gain and weight loss, the body weight and fat mass of WC animals stabilized at a higher steady-state level than the LFD-lean controls (**Figure 1A-B**). Although average energy intake during the entire weight loss phase was not different between WC and LFD-lean mice, WC mice exhibited significantly higher energy intake during the weight-stable phase (week 19-29), while feed efficiency was still reduced (**Figure 1G-H**). However, re-exposure of 10-week weight-stable WC mice to HFD feeding for 10 weeks increased fat mass to a level similar to the HFD-obese controls (**Figure 1A-B**).

### Impact of WC during leanness: Weight cycled-lean mice display WAT dysfunction despite normalized whole-body metabolic homeostasis

In order to investigate manifestations of obesogenic memory after prolonged weight stabilization, half of the animals were sacrificed after 29 weeks when WC mice were weight stable for 10 weeks (WC-lean), and compared with LFD-lean control animals (**Figure 2A**). Indeed, body weight and fat mass were higher in WC-lean mice compared to the LFD-lean controls, likely due to higher HFD exposure and cumulative energy intake (**Figure 2B-E**). While liver weights were fully normalized in WC-lean mice, the subcutaneous, inguinal WAT (iWAT) depot remained heavier compared to the LFD-lean mice. Weights of visceral, epididymal white adipose tissue (eWAT) were similar across all groups (**Figure 2F**), congruent with previous work using prolonged HFD exposures (33).

**Figure 2:**
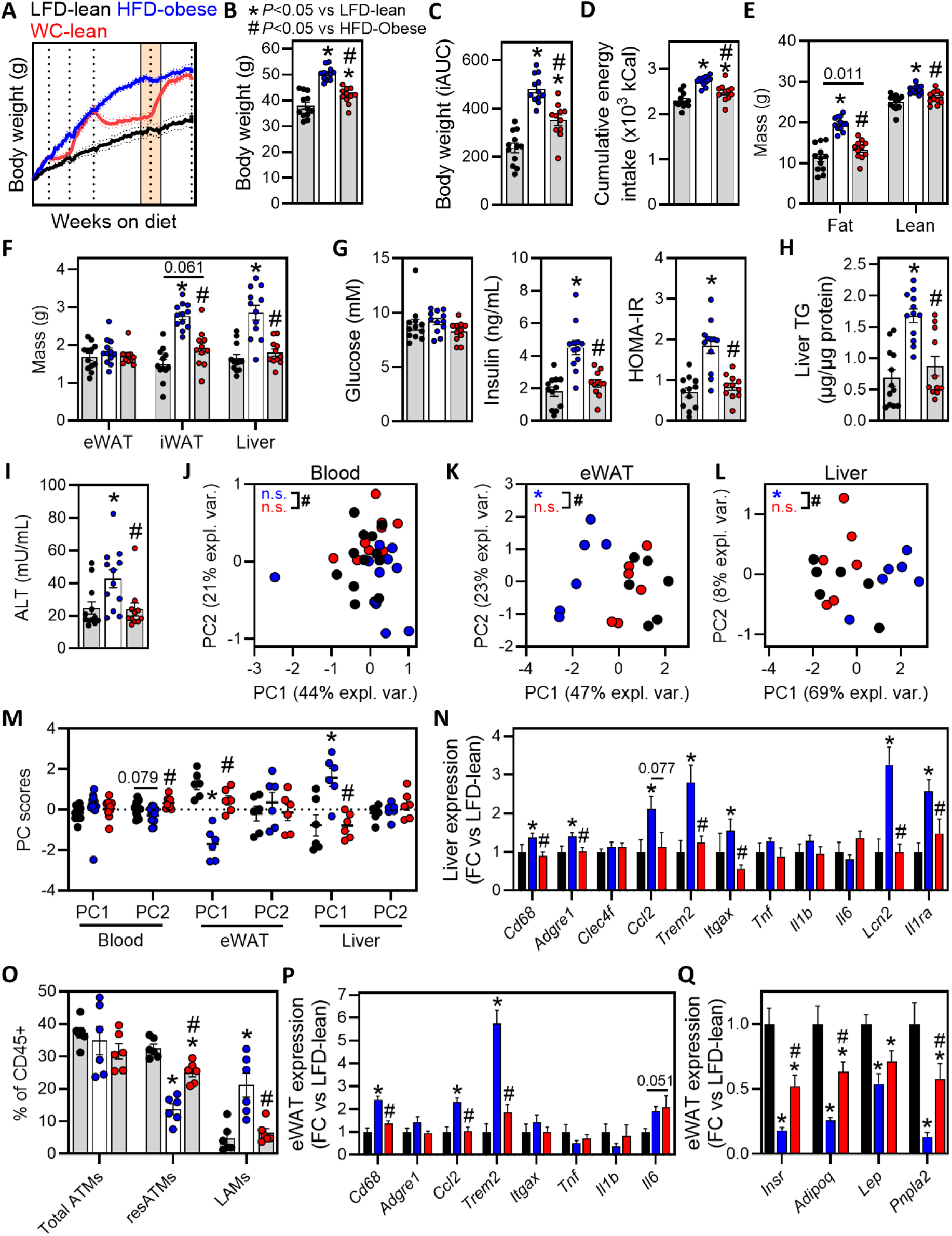
Weight cycled-lean mice display features of eWAT dysfunction despite normalized whole-body metabolic homeostasis. (**A**) Half of the LFD-lean (black), HFD-obese (blue) and weight-cycled-lean (WC-lean, red) mice were sacrificed when WC-lean were weight stable for 10 weeks. (**B**) Body weight measured at sacrifice. (**C**) Incremental area under the curve of body weight. (**D**) Cumulative energy intake throughout the experiment. (**E**) Fat and lean mass determined at sacrifice. (**F**) Weights of visceral epidydimal white adipose tissue (eWAT), subcutaneous inguinal WAT (iWAT) and liver determined at sacrifice. (**G**) Fasting blood glucose and plasma insulin were used to calculate HOMA-IR at 1 week before sacrifice. (**H**) Liver triglyceride levels were determined as a proxy for hepatic steatosis. (**I**) Plasma ALT levels were determined as a proxy for liver injury. (**J-L**) Immune cells were isolated from blood, eWAT and liver and phenotyped by flow cytometry. Full gating strategies are shown in Figure S1. Frequencies of immune cell subsets were used as input for unsupervised principal component analysis to assess variation between mice in the immune landscape in blood (J), eWAT (K) and liver (L). (**M**) Scores for the first two principal components per tissue. (**N**) Liver gene expression of the indicated inflammatory markers. (**O**) Total adipose tissue macrophages (ATMs), resident ATMs and lipid-associated ATMs as frequencies of CD45+ cells in eWAT determined via flow cytometry. (**P-Q**) eWAT gene expression of the indicated inflammatory markers (P) and markers of adipose tissue function (Q). Data are expressed as mean ± SEM (n=11-12 mice per group for B-J, M-N and P-Q; n=6 pools of 1-2 mice per group for K-M and O). Significance was tested by One-Way ANOVA with Tukey correction for multiple testing (B-I and M-Q) or pair-wise PERMANOVA (J-L). * *p* < 0.05 vs LFD-lean, # *p* < 0.05 vs HFD-obese.

Despite higher body weight and fat mass, obesity-induced changes in fasting plasma total cholesterol (TC), non-esterified fatty acids (NEFA), insulin and HOMA-IR, as proxies for insulin resistance, all completely normalized after WC-lean mice were weight stable for 10 weeks (**Figure S2A**, **Figure 2G**). Moreover, liver triglycerides (TG) and plasma transaminase (ALT) levels were also normalized (**Figure 2H-I**), indicating a complete reversal of MASLD.

To determine the impact of WC on obesity-induced chronic low-grade inflammation, we immunophenotyped blood, eWAT and liver via flow cytometry (see **Figure S1** for gating strategies and all detected immune cell types). Of note, the numbers of leukocytes per mL blood or per g tissue were not significantly different between groups (**Figure S2B-D**). We used principal component analysis (PCA) followed by PERMANOVA on immune cell composition data to evaluate whether immune landscapes were different, and which immune cell subsets contributed to these differences. Immune cell composition was significantly different in eWAT and liver of HFD-obese control compared with LFD-lean control animals, and tended to be different in blood. This was explained by PC2 in blood, and PC1 in both the eWAT and liver PCA (**Figure 2J-M**). Evaluating the top 5 immune cell subsets that contributed most to these PC scores revealed both myeloid and lymphoid immune cell subsets (**Figure S2E-G**). Notably, eWAT variation along PC1 was explained by lipid-associated macrophages (LAMs), eosinophils, resident adipose tissue macrophages (resATMs) and CD8 T cell subsets, which were indeed markedly changed in HFD-obese compared with LFD-lean controls (**Figure S2F**), as expected (14, 34). In the liver, the top 5 contributing features to PC1 were monocyte-derived macrophages (moMacs), monocyte-derived Kupffer cells (moKCs), and effector CD8 T cells, NK T cells and naïve CD4 T cells, which were differentially regulated by obesity (**Figure S2G**) (14, 20, 21, 34).

Importantly, the liver immune landscape completely normalized in WC-lean mice (**Figure 2L-M, Figure S2G**). Moreover, expression of obesity-induced inflammatory hepatic gene markers, encompassing markers of Macrophages/Kupffer cells and their recruitment (*Cd68, Adgre1, Clec4f, Ccl2*), markers of lipid-associated macrophages (*Trem2, Itgax*), and proinflammatory cytokines and their receptors (*Tnf, Il1b, Il6, Lcn2, Il1ra*) also completely reverted to the levels observed in LFD-lean mice (**Figure 2N**). In addition, plasma proteomics using a panel of 48 inflammatory markers revealed no distinct clustering of the plasma inflammatory proteomes of LFD-lean and WC-lean animals (**Figure S2H-I**). In contrast, eWAT retained some obesity-related inflammatory features despite long-term weight stabilization, as resATMs remained significantly decreased and eWAT *Il6* expression tended to be higher in WC-lean animals compared with the LFD-lean controls (**Figure 2K,M,O,P**). In addition, expression of adipose tissue functional genes, i.e. *Insr* (encoding Insulin Receptor), *Adipoq* (encoding Adiponectin), *Lep* (encoding Leptin) and *Pnpla2* (encoding Adipose Triglyceride Lipase [ATGL]), remained significantly downregulated in eWAT of WC-lean mice (**Figure 2Q**), suggesting a lasting imprint of WC on eWAT function.

The apparent body weight and adiposity set- or settling-point alterations induced in WC-lean mice prompted us to assess the impact of WC on hypothalamic inflammation, as the arcuate nucleus (ARC) of the hypothalamus is a central regulator of body weight (via energy intake and expenditure (26, 27)), and ARC microglia have been shown to play a critical role in regulating these processes (35). Confocal microscopy of hypothalamic sections (**Figure 3A**) revealed a trend towards a reduced total number of Iba1+ microglia in WC-lean animals as compared to LFD-lean controls. Notably, nuclear-pNFκB/Iba-1 double-positive microglia were 60% less abundant in WC-lean mice (**Figure 3B-C**). Interestingly, 3D reconstruction revealed that the mean cell volume of nuclear-pNFκB/Iba-1 double-positive microglia was significantly increased in WC-lean mice (**Figure 3D-E**), correlating with lower microglia numbers (**Figure S3**, **Figure 3F**). Jointly considering nuclear-pNFκB/Iba-1 double-positive microglia in the two groups of lean mice demonstrated that activated ARC microglia were distinct, being less abundant and larger in the WC-lean group (**Figure 3F**). Moreover, hypothalamic-ARC microglia counts and mean cellular volume correlated with metabolic parameters in the two lean groups combined (**Figure S3**). Both a lower number and higher volume of nuclear-pNFκB/Iba-1 double-positive ARC microglia significantly correlated with higher fat mass (**Figure 3G**). In addition, fewer microglia (as in WC-lean mice) correlated with lower feed efficiency but higher adiposity, whereas a larger activated microglia volume correlated with higher adiposity and fasting insulin (**Figure S3** and **Figure 3G**).

**Figure 3:**
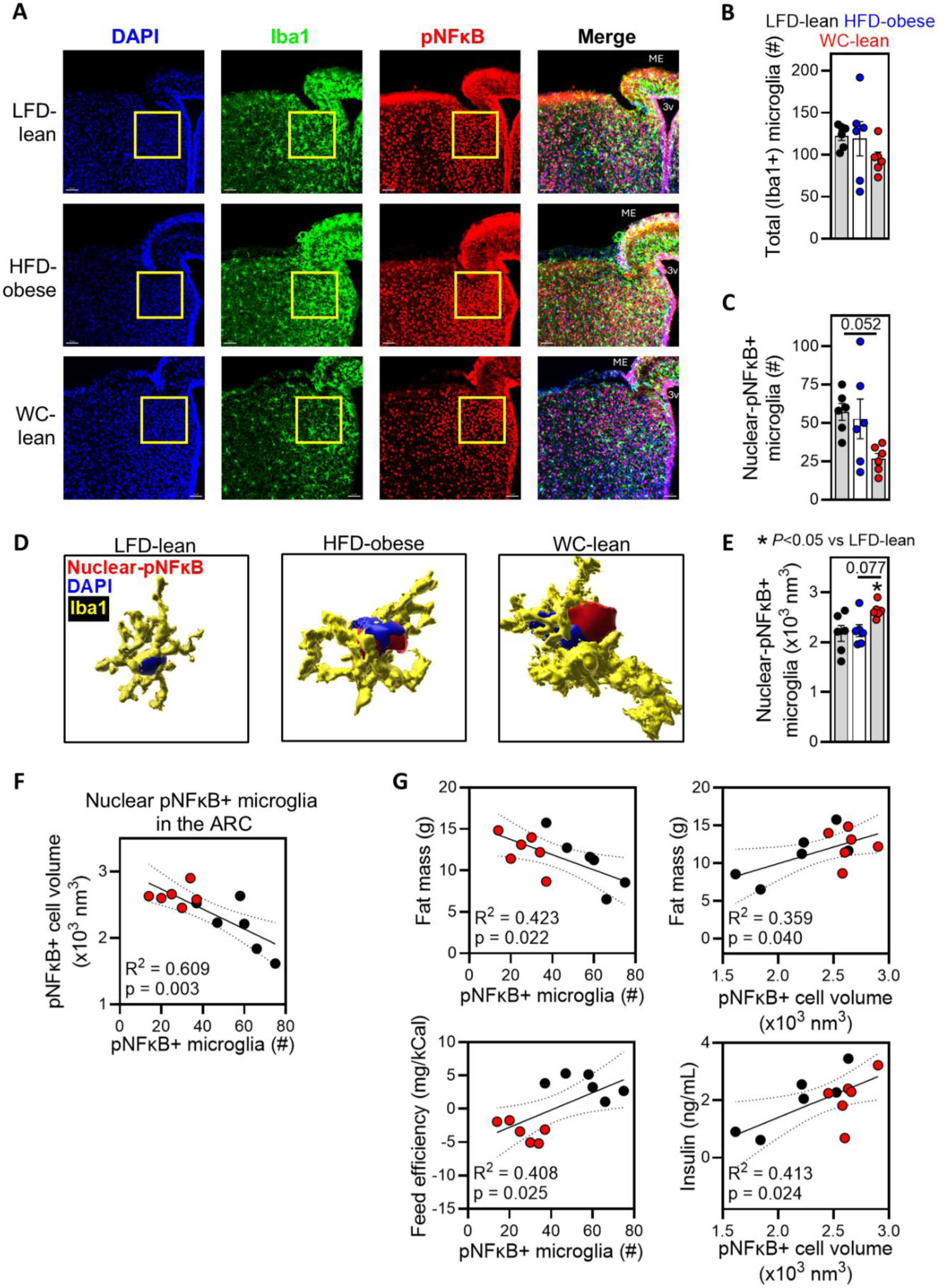
Adiposity and insulin resistance correlate with weight cycling-induced microglial changes in the arcuate nucleus of the hypothalamus. (**A**) Mice were treated and sacrificed as described in the legend of Figure 2. Brains were collected and fixed at sacrifice and cryosections of the hypothalamus were stained for nuclei (DAPI), microglia (Iba1) and phospho-NFκB (pNFκB). Scale bar indicates 50 μm. ME = median eminence, 3V = third ventricle. The yellow rectangle indicates the analyzed area. (**B-C**) Quantification of numbers of microglia and pNFκB+ inflammatory microglia in the arcuate nucleus of the hypothalamus. (**D-E**) Analysis of nuclear-pNFκB/Iba1 double-positive microglial cell volume. Representative image (D) and quantification (E). (**F-G**) Correlations between indicated parameters in WC-lean and LFD-lean control mice. Data are expressed as mean ± SEM or ±95% confidence interval (n=6 mice per group). Significance was tested by One-Way ANOVA with Tukey correction for multiple testing (B, C, E) or F-test for the regression model (F-G). * *p* < 0.05 vs LFD-lean.

Taken together, while obesity-induced insulin resistance and MASLD were fully resolved, eWAT exhibited a subtle inflammatory imprint of WC in long-term leanness, which was associated with fewer yet larger activated ARC microglia.

### Impact of WC in obesity: Weight cycling did not aggravate obesity-associated metabolic dysfunctions upon weight regain, but altered immune landscapes in metabolic organs

To determine whether a history of WC exacerbates metabolic tissue inflammation and whole-body insulin resistance in response to a subsequent weight regain, WC mice were re-exposed to HFD for an additional 10 weeks (WC-obese) and compared with HFD-obese control animals (**Figure 4A**). Interestingly, body weight, cumulative energy intake and body composition were similar between WC-obese and HFD-obese mice, despite lower cumulative HFD exposure in WC-obese mice (**Figure 4B-F**). Moreover, no changes were observed in fasting TG, TC and NEFA levels between WC-obese and HFD-obese animals (**Figure S4A**). In contrast to our expectations, WC did not increase fasting glucose, fasting insulin and HOMA-IR compared with HFD-obese controls (**Figure 4G**), indicating no difference in whole-body insulin resistance. An oral glucose tolerance test additionally revealed that WC did not aggravate obesity-induced glucose intolerance (**Figure 4H-I**). Moreover, plasma insulin concentration at t=20 minutes post glucose gavage was similar between HFD-obese and WC-obese mice (**Figure 4J**), suggesting no difference in glucose-stimulated insulin secretion.

**Figure 4:**
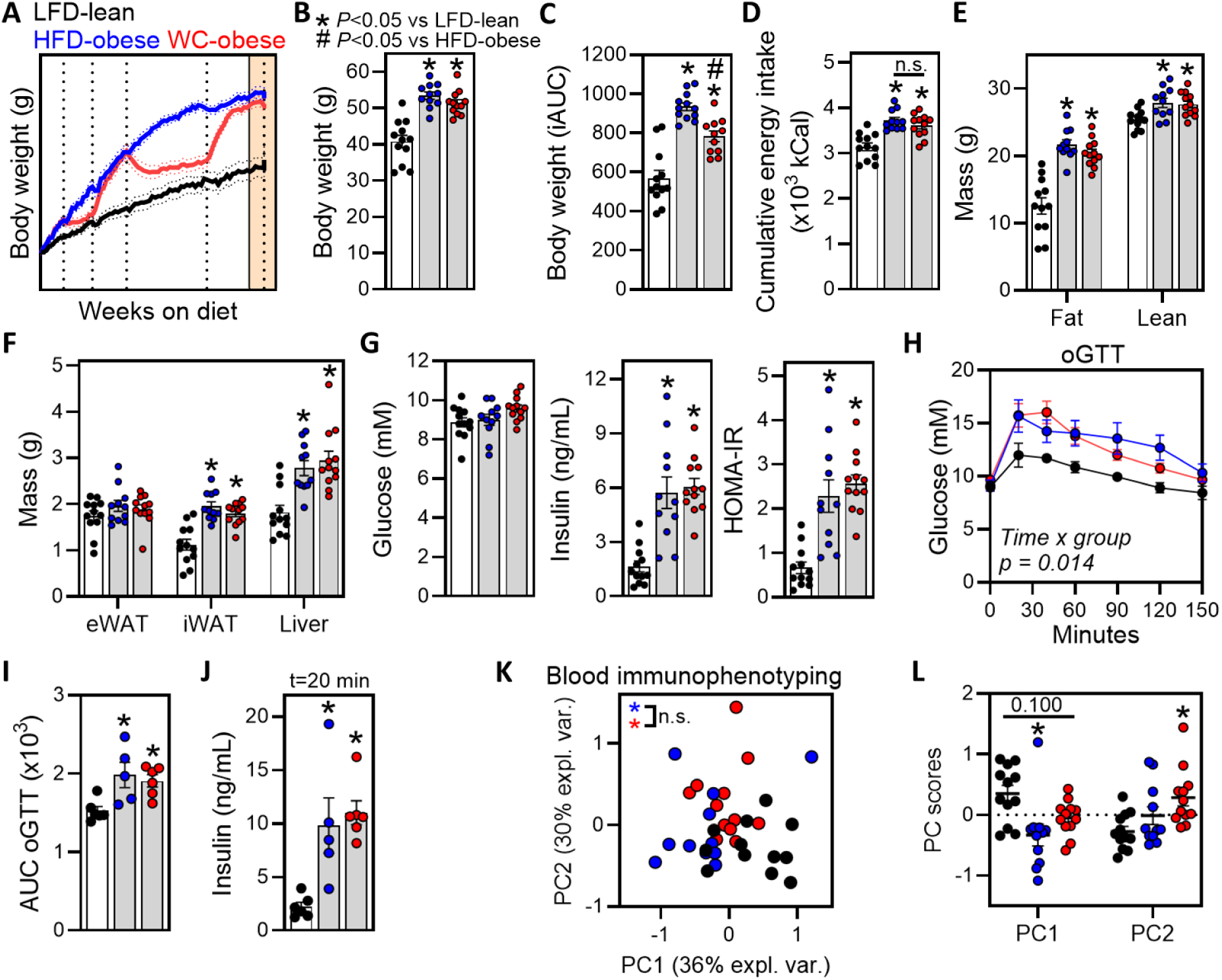
Weight cycling did not aggravate obesity-associated metabolic dysfunctions upon weight regain. (**A**) LFD-lean (black), HFD-obese (blue) and weight-cycled-obese (WC-obese, red) mice were sacrificed after a 10-week re-exposure of WC mice to HFD. (**B**) Body weight measured at sacrifice. (**C**) Incremental area under the curve of body weight. (**D**) Cumulative energy intake throughout the experiment. (**E**) Fat and lean mass determined at sacrifice. (**F**) Weights of eWAT, iWAT and liver were determined at sacrifice. (**G**) Fasting blood glucose and plasma insulin were used to calculate HOMA-IR at 1 week before sacrifice. (**H-I**) An oral glucose tolerance test was performed at 1 week before sacrifice. Glucose excursion curve (**H**) and area under the curve for glucose (I) are shown. (**J**) Plasma insulin was measured at 20 minutes post glucose gavage. (**K**) Principal component analysis to assess variation in the blood immune cell composition as described in the legend of Figure 2. (**L**) Scores for the first two principal components. Data are expressed as mean ± SEM (n=11-12 mice per group for B-G and K-L; n=5-6 mice per group for H-J). Significance was tested by One-Way ANOVA with Tukey correction for multiple testing (B-G, I-J and L), Two-Way ANOVA with Tukey correction for multiple testing (H) or pair-wise PERMANOVA (K). * = *p* < 0.05 vs LFD-lean, # = *p* < 0.05 vs HFD-obese.

WC did not impact circulating immune cell numbers upon weight regain (**Figure S4B**). Additionally, while HFD-obese and WC-obese animals clustered significantly differently from LFD-lean control animals via PCA on circulating immune cell composition, which was mostly driven by PC1. Circulating immune cell composition was largely similar in HFD-obese controls and WC-obese mice, with subtle changes in circulating T cell subsets (**Figure 4K-L**, **Figure S4C**). Finally, plasma proteomics revealed small differences in the inflammatory proteome between HFD-obese and WC-obese mice, where IL-5, CTLA4 and IL-9 were reduced, and IL-12a/b was increased, although absolute values were low (<1 pg/mL, data not shown) (**Figure S4D-E**).

We next investigated the impact of WC on the liver after weight regain. WC did not further increase the obesity-induced intracellular lipid droplet area, hepatic steatosis, liver TG and plasma ALT levels compared with HFD-obese control mice (**Figure 5A-E**). Moreover, gene expression of the fatty acid transporter CD36, the lipid droplet-associated protein CIDEA, the rate-limiting enzyme of fatty acid oxidation CPT1A, and markers of hepatic fibrosis and inflammation were similar between WC-obese and HFD-obese control mice (**Figure S5A-B**, **Figure 5F**).

**Figure 5:**
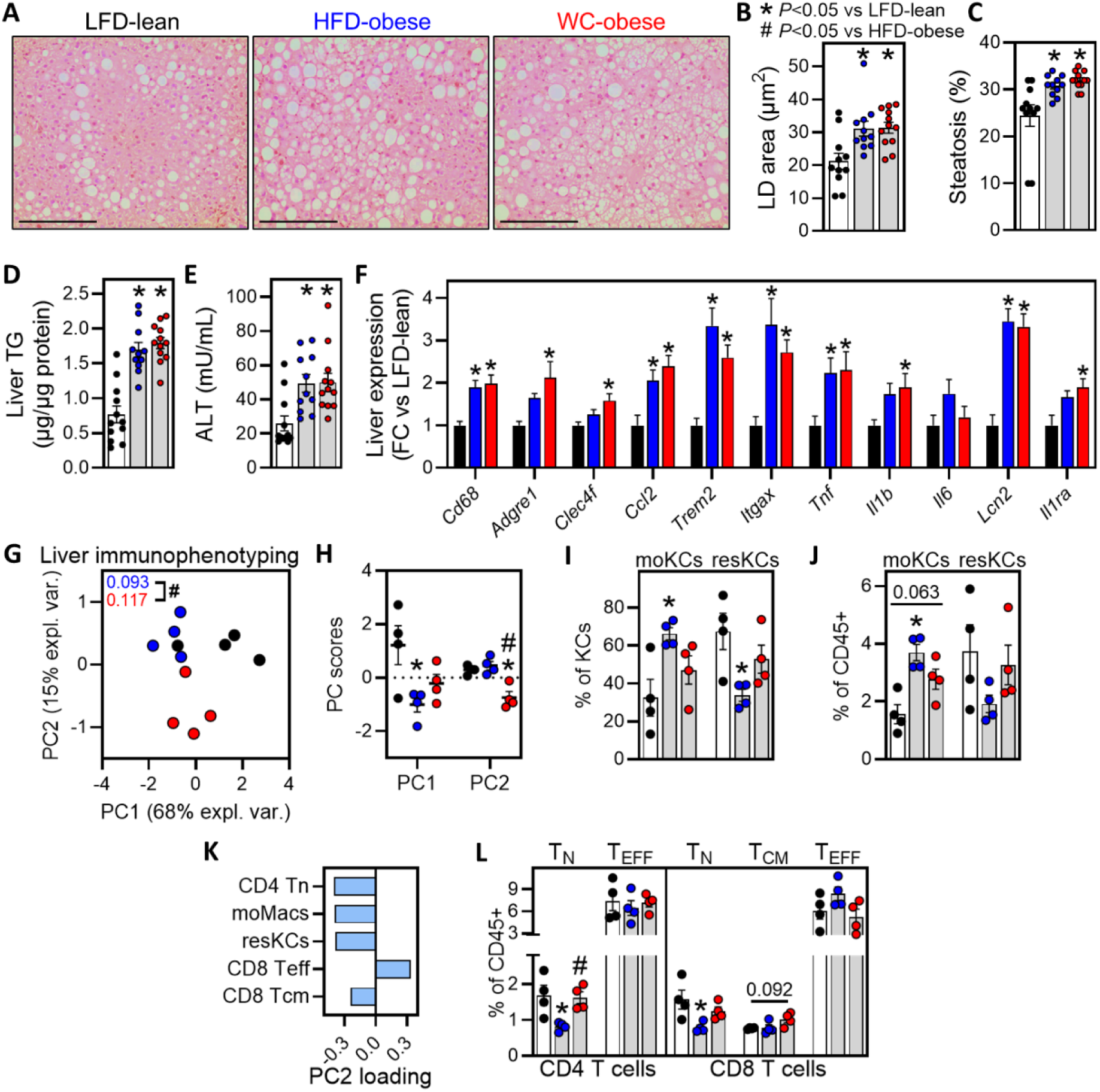
Weight cycling did not aggravate hepatic steatosis upon weight regain, but modified the liver immune landscape. (**A**) LFD-lean (black), HFD-obese (blue) and weight-cycled-obese (WC-obese, red) mice were sacrificed after a 10-week re-exposure of WC mice to HFD. A part of the liver was sectioned and H&E stained. (**B-C**) Average lipid droplet area (B) and percentage of steatosis (C) were determined from H&E-stained slides. (**D-E**) Liver triglycerides (D) and plasma ALT levels (E) were determined. (**F**) Hepatic gene expression of the indicated inflammatory markers. (**G**) Principal component analysis to assess variation in the liver immune cell composition as described in the legend of Figure 2. (**H**) Scores for the first two principal components. (**I-J**) Frequencies of indicated hepatic myeloid subsets determined via flow cytometry. (**K**) Top 5 hepatic immune cell subsets contributing to the PC2 score. (**L**) Frequencies of indicated hepatic T cell subsets. Data are expressed as mean ± SEM (n=11-12 mice per group for B-F; n=4 pools of 1-6 mice per group for G-L). Significance was tested by One-Way ANOVA with Tukey correction for multiple testing (B-F, H-J and L) or pair-wise PERMANOVA (G). * = *p* < 0.05 vs LFD-lean, # = *p* < 0.05 vs HFD-obese.

We next assessed the liver immune cell composition to further dissect hepatic inflammation. While the number of leukocytes per g liver was similar between groups (**Figure S5C**), unsupervised PCA revealed distinct clustering of all groups. Indeed, HFD-obese and WC-obese groups tended to be differentially clustered from LFD-lean controls, which was mostly explained by PC1 (**Figure 5G-H**), where myeloid cells strongly contributed to the variance (**Figure S5D**). Indeed, hepatic monocytes and moMacs that develop into moKCs (20, 21) were more abundant in both HFD-obese and WC-obese mice compared with LFD-lean controls (**Figure S5D-E, Figure 5I-J**). Interestingly, while hepatic monocytes and moMacs were similar between WC-obese and HFD-obese controls, moKC and resKC abundance did not reach HFD-obese levels in WC-obese animals (**Figure 5I-J**). Indeed, the WC-obese group clustered significantly differently from HFD-obese controls on PC2 (**Figure 5G-H**), at least partly explained by naïve CD4 T cells, moMacs, resKCs, and effector and central memory CD8 T cells (**Figure 5K-L**), of which the abundance was more similar to the LFD-lean control group. Of note, cell size and granularity of moKCs and resKCs, and expression of the lipid-associated macrophage markers CD9 and CD11c were not significantly different between groups (**Figure S5F-G**). Altogether, these data indicate that the liver immune landscape of WC-obese mice incompletely reverted to HFD-obese controls upon weight regain, which coincided with no aggravation of hepatic steatosis, hepatic fibrosis and expression of inflammatory genes.

WC subjects WAT to repetitive expansion and contraction, which has been reported to induce cellular stress and inflammation (36). Thus, we investigated the impact of WC on visceral eWAT upon weight regain. Adipocyte size and expression of adipose tissue functional markers were similar between HFD-obese and WC-obese mice (**Figure 6A-C**), indicating that WC did not significantly aggravate adipose tissue dysfunction. In addition, crown-like structures of macrophages surrounding adipocytes, a hallmark of obese adipose tissue, were similarly increased in both HFD-obese and WC-obese animals (**Figure 6D**), and WC did not result in increased eWAT inflammatory gene expression in comparison with HFD-obese mice (**Figure 6E**).

**Figure 6:**
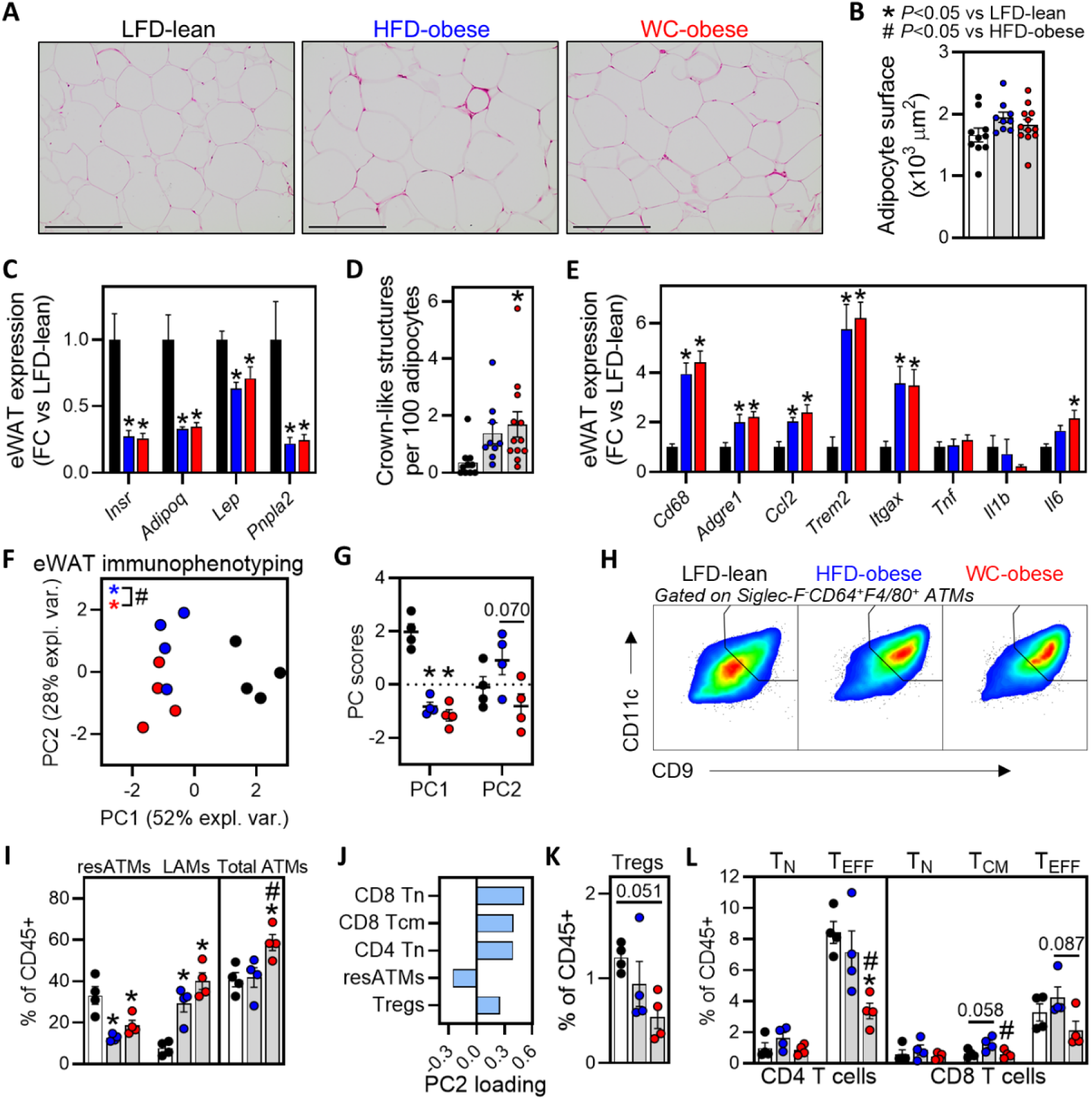
Weight cycling increased adipose tissue macrophages and reduced T cells upon weight regain. (**A**) LFD-lean (black), HFD-obese (blue) and weight-cycled-obese (WC-obese, red) mice were sacrificed after a 10-week re-exposure of WC mice to HFD. eWAT was sectioned and H&E stained. (**B**) Average adipocyte surface area was determined from H&E-stained slides. (**C**) eWAT gene expression of the indicated markers of adipose tissue function. (**D**) Crown-like structures were quantified from H&E-stained slides. (**E**) eWAT gene expression of the indicated inflammatory markers. (**F**) Principal component analysis to assess variation in the eWAT immune cell composition as described in the legend of Figure 2. (**G**) Scores for the first two principal components. (**H**) Representative gating of lipid-associated macrophages (LAMs). resATMs were determined as not-LAMs. (**I**) Frequencies of indicated eWAT macrophage subsets. (**J**) Top 5 eWAT immune cell subsets contributing to the PC2 score. (**K-L**) Frequencies of indicated eWAT T cell subsets. Data are expressed as mean ± SEM (n=11-12 mice per group for B-E; n=4 pools of 1-6 mice per group for F-L). Significance was tested by One-Way ANOVA with Tukey correction for multiple testing (B-E, G, I, K-L) or pair-wise PERMANOVA (F). * = *p* < 0.05 vs LFD-lean, # = *p* < 0.05 vs HFD-obese.

However, while leukocytes per g eWAT were similar between HFD-obese and WC-obese mice (**Figure S6A**), all groups clustered significantly differently via PCA on immune cell composition (**Figure 6F-G**). Indeed, while PC1 captured the separation of obese from lean animals, among others explained by eosinophils, ILC2s, resATMs and LAMs, PC2 separated HFD-obese from WC-obese mice (**Figure S6B-C**). Interestingly, WC-obese mice did not exhibit an increase in eWAT B cells similar to HFD-obese controls, while total ATMs were increased (**Figure S6C**, **Figure 6H-I**). This increase in ATMs was associated with a significant upregulation of *Ccl2* in eWAT adipocytes separated from the stromal vascular fraction (**Figure S6D-G**). Of note, the expression of LAM markers (e.g. CD9, CD11c and lipidTOX) was similar between HFD-obese and WC-obese animals (**Figure S6H-I**). In addition to macrophages, T cell subsets also contributed to the separation of WC-obese from HFD-obese mice on PC2 (**Figure 6J**). Indeed, CD4 and CD8 T cell subsets were markedly reduced in WC-obese compared with HFD-obese control eWAT (**Figure 6K-L**).

To investigate the impact of WC on adipocyte *Ccl2* expression, we developed an in vitro model of adipose tissue weight cycling. Here, we switched 3T3-L1 adipocytes between high (25 mM) and low (1 mM) glucose concentrations in 3-day intervals for a total of 12 days (**Figure S6J-L**). Strikingly, in vitro WC in 3T3-L1 adipocytes increased *Ccl2* expression and resulted in increased MCP-1 production in the supernatant (**Figure S6M-N**), suggesting that WC may increase adipocyte-mediated macrophage recruitment to visceral adipose tissue. Altogether, while immune landscapes in eWAT and liver were modified, with increased adipose tissue macrophage recruitment and reduced effector T cells, WC did not aggravate obesity-induced insulin resistance and MASLD upon weight regain.

## Discussion

Body weight cycling, or yo-yo dieting, is a prevalent phenomenon following intentional weight loss (9) and is suggested to exacerbate obesity-associated complications (12, 13). Here, we show that animals with a history of WC stabilize at a higher body weight following weight loss, despite normalization of whole-body metabolic homeostasis and metabolic tissue inflammation. These changes were associated with lasting features of eWAT dysfunction and fewer but larger inflammatory microglia in the hypothalamus. Finally, although WC reprogrammed immune landscapes in eWAT and liver upon weight regain, characterized by increased eWAT macrophages and reduced eWAT and liver effector T cells, these inflammatory changes were uncoupled from whole-body metabolic dysfunctions.

We demonstrate that prolonged weight stabilization after weight loss via *ad libitum* LFD feeding completely normalized markers of obesity-induced insulin resistance. In addition, MASLD and liver inflammation also fully reversed to lean control levels, underscoring the unique regenerative capacity of the liver (37). Others have also reported normalization of metabolic homeostasis following weight loss (25, 38), yet few also investigated liver inflammation. In fact, an inflammatory hepatic gene signature was detected directly after weight loss (23), whereas inflammatory cytokine expression normalized after 8 weeks of weight stabilization (24). We additionally show that given sufficient time of weight stabilization in leanness, liver immune cell composition completely normalizes, including the normalization of proinflammatory, obesity-induced moMacs and moKCs, and reconstitution of the resKC niche that is lost in the steatotic liver (20, 21). resKCs are seeded during embryogenesis, whereas moKCs are bone marrow monocyte-derived. This difference in macrophage ontogeny has been identified as an important factor determining macrophage activation (39), as reflected by the increased inflammatory potential of moKCs (20). Both moKCs and resKCs have recently been reported to contribute to tissue repair and revert to baseline levels once the injury is resolved (40). However, whether the remaining embryonically-seeded resKCs proliferate to replenish the resKC niche, or whether the monocyte-derived cells convert into tissue-resident macrophages as described in other compartments (41), remains to be determined. It would thus seem important to assess whether converted resKCs respond differently to future inflammatory triggers, such as re-exposure to MASLD, tissue toxicity or cancer.

Although metabolic homeostasis and liver inflammation normalized, fat mass remained elevated and features of eWAT dysfunction persisted in WC-lean mice. eWAT has been demonstrated to retain an inflammatory fingerprint following weight loss, both acutely and after weight stabilization (23, 24, 38). In particular, increased macrophage proliferation that maintains a LAM-like phenotype following weight loss was suggested to feed adipose tissue inflammation (25). In our study, LAMs abundance normalized following long-term weight stabilization on an LFD in WC-lean mice, while resATM abundance was lower compared with LFD-lean controls. Since eWAT *Il6* expression remained elevated and WAT functional genes downregulated, together this suggests that WC induces a subtle imprint on eWAT even after long-term weight stabilization, which is a potentially lasting manifestation of obesogenic memory. One may hypothesize this imprint contributes to accelerated weight regain (42), or to increased susceptibility to secondary inflammatory triggers (43).

It is worth considering whether this incomplete restoration of WAT homeostasis in WC-lean mice, compared with LFD-lean controls, is secondary to elevated body weight and adiposity. Weight loss and gain acutely impact metabolic dysfunctions, improving or worsening it, respectively, which results from changes in energy balance. To control for this, we used an experimental design in which WC mice reached a steady-state body weight after weight loss and gain. Interestingly, WC mice stabilized on higher body weight after two weight gain and weight loss phases under *ad libitum* LFD feeding, which implies that body weight regulation is impaired. This was associated with microglial adaptations in the ARC of the hypothalamus, the central regulator of energy homeostasis (26, 27). Mechanistically, others have shown that HFD-induced gut microbial changes contribute to the rate of post-dieting weight regain. These changes increased intestinal lipid absorption and resulted in decreased energy expenditure (28, 44). In addition, a gut-brain axis via vagal nerve signaling and/or production of intestinal endocrine factors has been demonstrated to impact satiety and energy expenditure (45). Such mechanisms could have contributed to WC-induced elevated weight stabilization in our experimental settings, potentially mediated and/or reflected by microglial adaptations, although this requires further study. Alternatively, adipocytes were recently demonstrated to epigenetically memorize obesity following weight loss (42). One may thus speculate that upon long-term WC, adipocytes contribute to defending a novel body weight settling point, *e.g.* via regulating their metabolism, adipogenesis and/or paracrine signaling to the brain through leptin.

In our study, mice that underwent WC did not show impaired insulin resistance, glucose intolerance and MASLD compared to similarly obese mice that did not undergo WC. These results differ from previous reports suggesting that WC exacerbates glucose intolerance (38, 46). Several differences in experimental design may explain this discrepancy. First, we used an HFD with 45% instead of 60% of calories derived from fat, the latter resulting in faster weight gain and worse metabolic dysfunctions, potentially enhancing WC-induced detrimental effects on metabolic health. However, the 45% HFD we used here may better resemble the dietary macronutrient composition of human (Western) diets (47). Second, we compared obese mice that underwent WC with mice that were chronically obese, rather than with late-onset obese animals. Although cumulative energy intake was similar between the WC-obese and HFD-obese groups, the duration of HFD exposure was only half in the WC-obese mice. A recent meta-analysis confirmed that WC only worsened metabolic outcomes when compared with late-onset obese animals (48), when HFD exposure is similar. However, a recent systematic review of studies also concluded that WC is not associated with adverse effects on body weight, body composition and metabolic rate (49). Thus, while WC may have modest adverse effects on metabolic health in obesity, our data indicate that weight loss followed by weight regain does not exacerbate metabolic dysfunctions relative to chronic obesity.

While metabolic dysfunctions were similar, WC did modulate the immune landscapes in the liver and eWAT upon weight regain compared with HFD-obese control mice. The impact of one cycle of weight loss and weight regain on eWAT immune cell composition has been previously reported (38), yet the impact of multiple cycles, and whether the liver immune cell composition is affected, remained elusive. We identified that eWAT macrophages were increased in WC-obese mice upon weight regain, which was associated with WC-induced *Ccl2* expression in adipocytes. Interestingly, *in vitro* nutrient cycling of 3T3-L1 adipocytes recapitulated this increase in *Ccl2* expression and MCP-1 production, suggesting this may be a feature of adipocyte-intrinsic obesogenic memory. In addition, eWAT effector CD4 and CD8 T cells and hepatic effector CD8 T cells were reduced in WC-obese compared with HFD-obese animals. Concurrently, hepatic naïve T cell subsets were increased. These results may correspond with an enhanced obesity-induced T cell exhaustion phenotype, at least in eWAT (38, 50), with currently unknown metabolic consequences. However, CD4 T cell memory has also been described to accelerate post-dieting weight regain (51) and a recent pre-print suggests that T cell memory contributes to aggravated glucose intolerance induced by WC (52). Together, this indicates that the contribution of T cells to WC-induced body weight regulation and metabolic dysfunctions is complex and requires further study.

As a recent report demonstrated that WC may result in bone marrow adaptations that protect against weight regain (53), we cannot exclude the possibility that WC-induced reprogramming of immune cell composition in eWAT, liver and potentially also the bone marrow, are a compensatory mechanism to preserve tissue and metabolic homeostasis. At the same time, a history of obesity or Western diet feeding has also been shown to increase harmful inflammatory responses in the context of neuroinflammation and atherosclerosis, respectively (43, 54). Given that obesity-induced T cell exhaustion and NK cell dysfunctions impair anti-tumor immunity (55, 56), further study of the impact of WC on obesity-associated tumorigenesis is warranted. Thus, while metabolic dysfunctions were not affected in our experimental settings, it remains possible that WC results in immune priming, which may predispose to a secondary inflammatory triggers with potential detrimental effects.

Altogether, we show that a history of WC results in elevated body weight stabilization even after prolonged *ad libitum* LFD feeding, suggesting induction of a novel body weight settling point. This is associated with lasting features of eWAT dysfunction and hypothalamic microglial adaptations. In obesity, WC does not aggravate metabolic dysfunctions upon weight regain, but may predispose to inflammation-driven morbidities given increased eWAT macrophage recruitment and reduced metabolic tissue effector T cells.

## Acknowledgments

Funding was provided by the consortium “The right timing to prevent type 2 diabetes” (TIMED; funded by The Netherlands Organization for Health Research and Development (ZonMw) [459001021], Dutch Diabetes Research Foundation (Diabetes Fonds) [2019.11.101], the Canadian Institutes of Health Research (CIHR) [TNC-174963], Health-Holland [LSHM20107]) and by ZonMw [09120012010062]. A.R. was funded by an Israel Science Foundation grant (ISF-194/24). The funders had no role in the design, analysis and reporting of the study. This study was supported by the Centre for Small Animals, Centralized Facilities for Animal Research at Wageningen University and Research (Wageningen, The Netherlands) and we greatly appreciate the support of their biotechnicians. We would like to thank Shohreh Keshtkar (Wageningen University & Research, Wageningen, The Netherlands) for technical assistance.

## Data availability

All the necessary data to evaluate the conclusions in the paper are provided in the paper itself and/or the Supplementary Materials. Raw data is available from the corresponding author upon reasonable request.

