## Supplemental tables and figures for "Weight cycling-induced hypothalamic and metabolic tissue immune remodeling is uncoupled from metabolic dysfunctions"

**Supplementary Material**

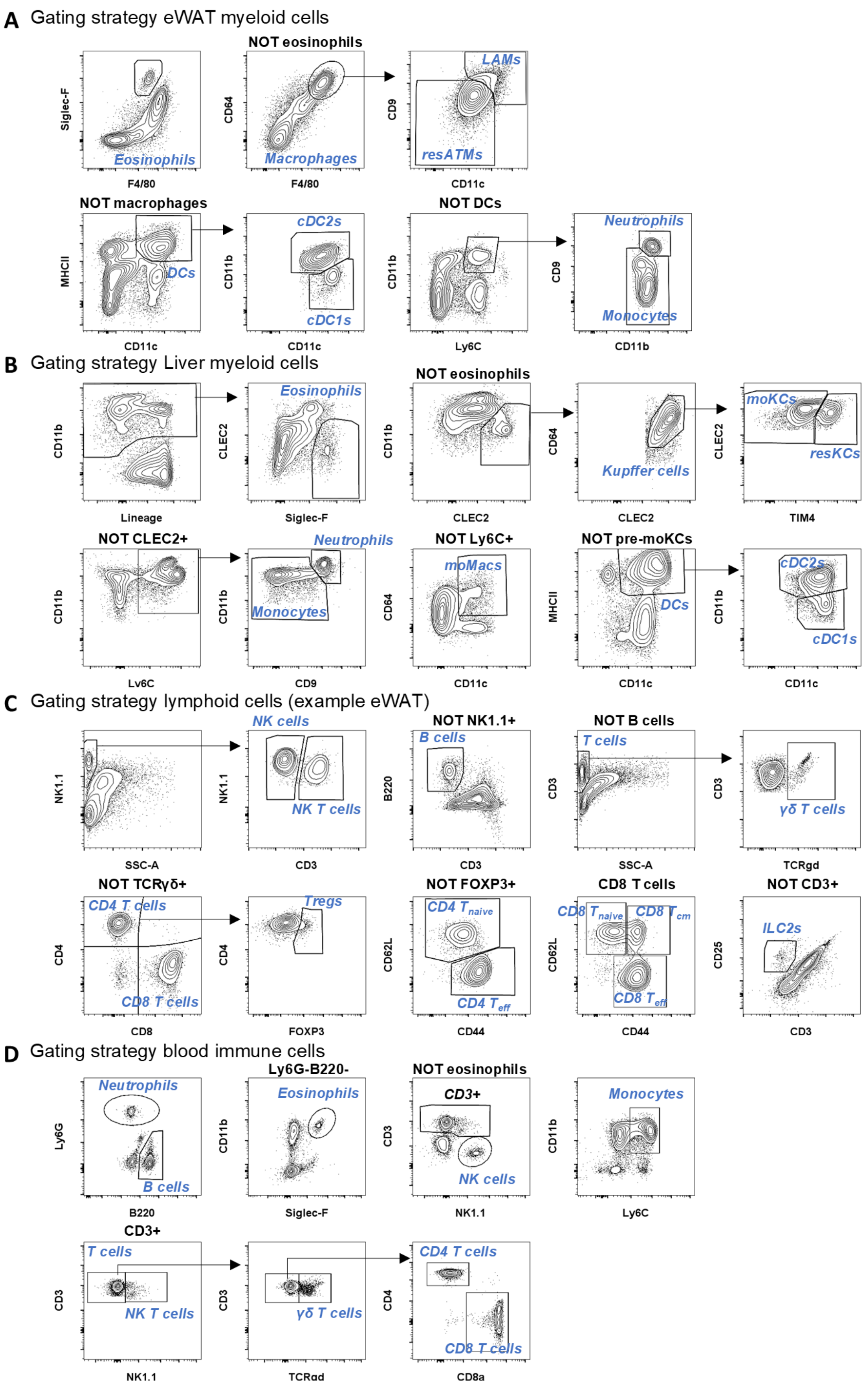


**Figure S1: Gating strategies**

(**A-D**) Representative gating strategies for indicated immune cell subsets in eWAT, liver and blood.

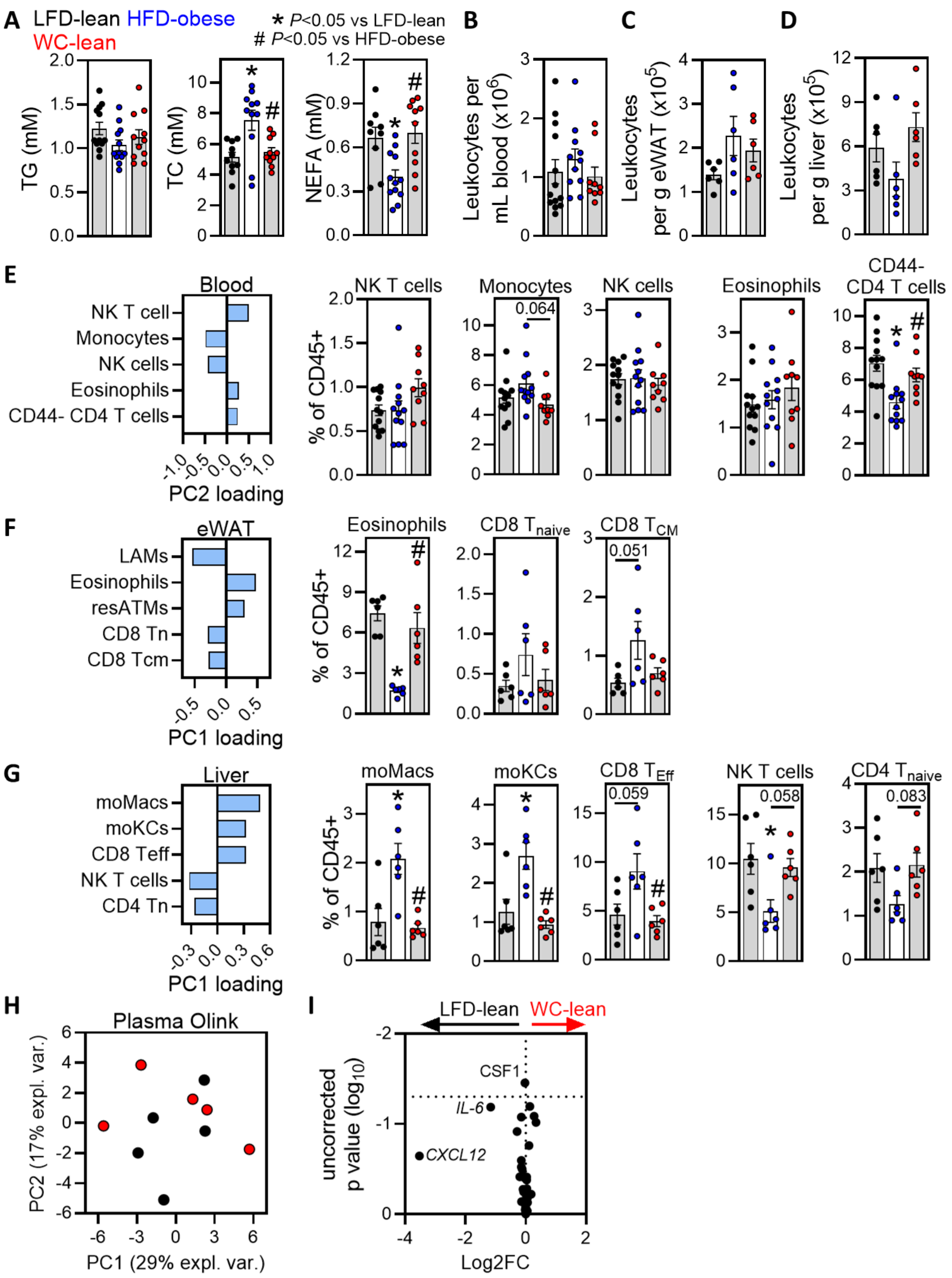


**Figure S2: Prolonged weight stabilization normalized immune cell landscapes in blood, epidydimal white adipose tissue and liver. Related to figure 2.**

(**A**) Half of the LFD-lean (black), HFD-obese (blue) and weight-cycled-lean (WC-lean, red) mice were sacrificed when WC mice were weight stable for 10 weeks. Fasting plasma triglycerides (TG), total cholesterol (TC) and non-esterified fatty acids (NEFA) were determined at 1 week before sacrifice. (**B-D**) Immune cells were isolated from blood (B), epidydimal white adipose tissue (eWAT; C) and liver (D) and numbers per mL or gram tissue were determined. (**E-G**) Top 5 blood (E), eWAT (F) and liver (G) immune cell subsets contributing to the PC1 score of the principal component analysis described in Figure 2, and frequencies of these immune cell subsets. (**H-I**) PCA (H) and volcano plot of $\log_2$ fold change and uncorrected p values (I) of plasma inflammatory markers determined via Olink comparing lean control and weight cycled mice. Data are expressed as mean ± SEM (n=9-12 mice per group for A-B and E; n=6 pools of 1-2 mice per group for C-D and F-G; n=5 mice per group for H-I). Significance was tested by One-Way ANOVA with Tukey correction for multiple testing (A-G) or independent samples t-test (I). * $p < 0.05$ vs LFD-lean, # $p < 0.05$ vs HFD-obese.

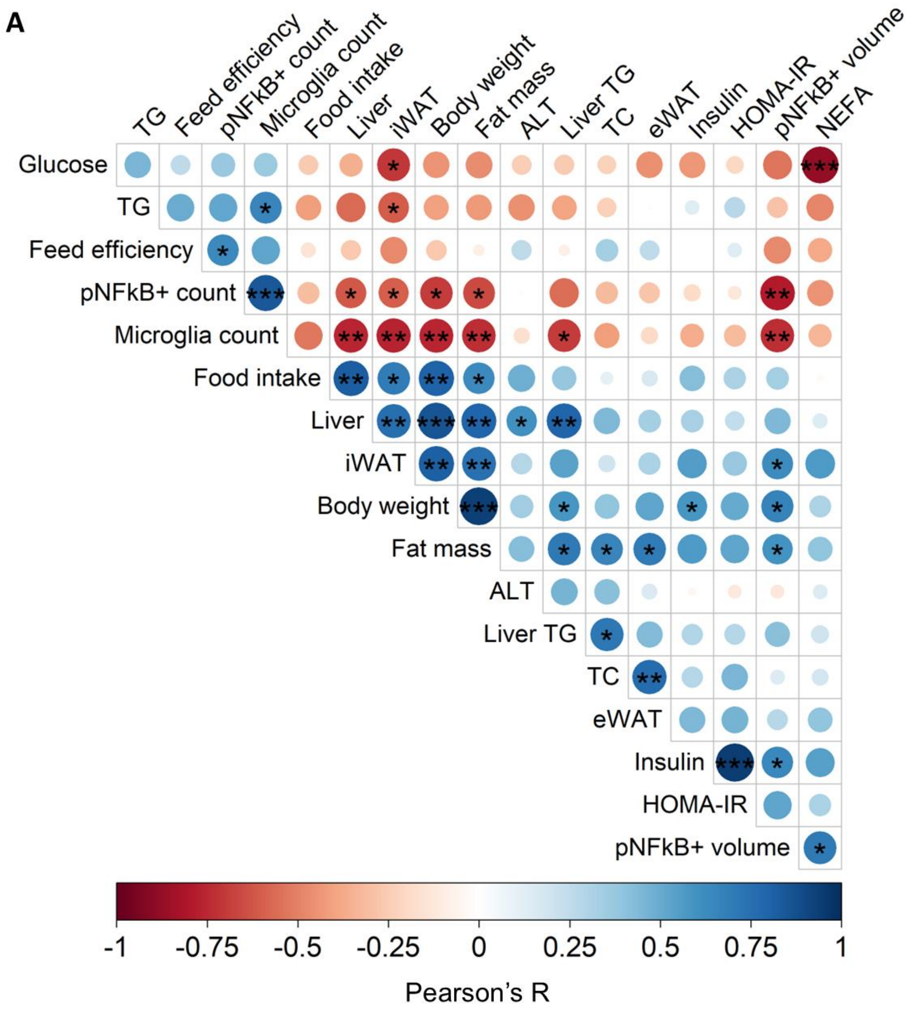


**Figure S3: Inflammatory hypothalamic microglia parameters correlated with feed efficiency, adiposity and insulin resistance. Related to figure 3.**

(**A**) Mice were treated and sacrificed as described in the legend of Figure 2. Correlogram for indicated microglial and metabolic parameters. Parameters were hierarchically clustered to indicate clusters of correlating parameters. Color intensity reflects Pearson's R values (n=6 mice per group). Significance was tested using a t-test for the correlation coefficient. * $p < 0.05$, ** $p < 0.01$, *** $p < 0.001$.

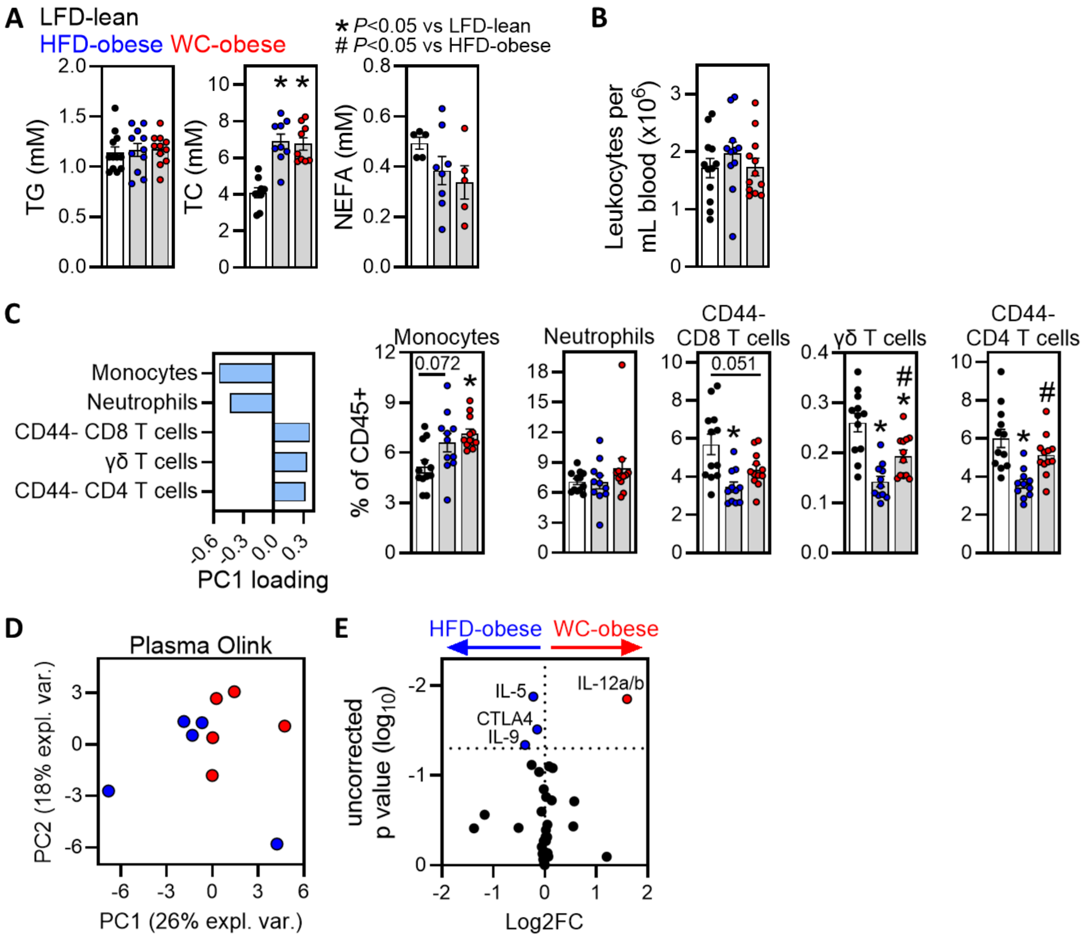


**Figure S4: Weight cycling marginally impacted circulating inflammatory parameters. Related to figure 4.**

(**A**) LFD-lean (black), HFD-obese (blue) and weight-cycled-obese (WC-obese, red) mice were sacrificed after a 10-week re-exposure of WC mice to HFD. Fasting plasma triglycerides (TG), total cholesterol (TC) and non-esterified fatty acids (NEFA) were determined at 1 week before sacrifice. (**B**) Immune cells were isolated from blood and numbers per mL blood were determined. (**C**) Top 5 blood immune cell subsets contributing to the PC1 score of the principal component analysis described in Figure 4, and frequencies of these immune cell subsets. (**D-E**) PCA (D) and volcano plot of $\log_2$ fold change and uncorrected p values (E) of plasma inflammatory markers determined via Olink comparing obese control and weight cycled mice. Data are expressed as mean ± SEM (n=5-12 mice per group for A-C; n=5 pools of 1-2 mice per group for D-E). Significance was tested by One-Way ANOVA with Tukey correction for multiple testing (A-C) or independent samples t-test (E). * $p < 0.05$ vs LFD-lean, # $p < 0.05$ vs HFD-obese.

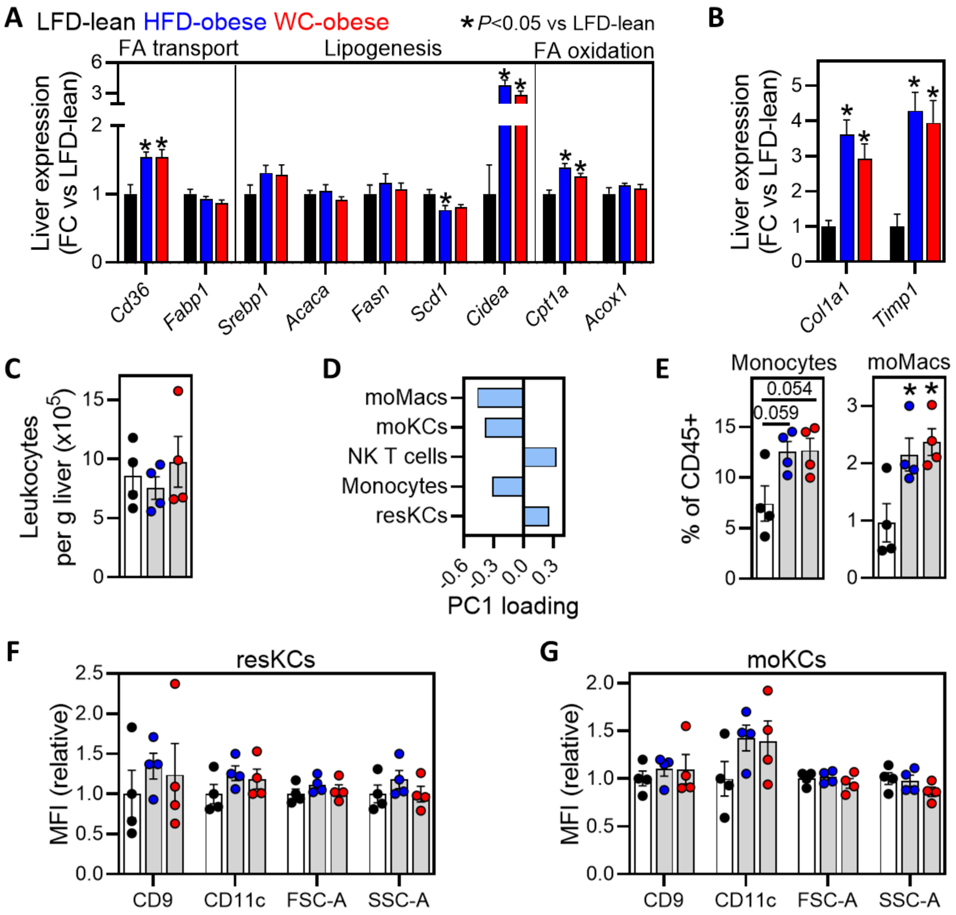


**Figure S5: Weight cycling neither aggravated hepatic steatosis nor affect expression of lipid-associated macrophage markers on Kupffer cells. Related to figure 5.**

(**A-B**) LFD-lean (black), HFD-obese (blue) and weight-cycled-obese (WC-obese, red) mice were sacrificed after a 10-week re-exposure of WC mice to HFD. Hepatic gene expression of the indicated lipid metabolism (A) and fibrosis (B) markers. (**C**) Immune cells were isolated from liver and numbers per g tissue were determined. (**D**) Top 5 hepatic immune cell subsets contributing to the PC1 score of the PCA on the hepatic immune cell composition. (**E**) Frequencies of indicated hepatic myeloid subsets determined via flow cytometry. (**F-G**) Quantification of expression of indicated markers on resident (F) and monocyte-derived (G) Kupffer cells determined via flow cytometry. Data are expressed as mean ± SEM (n=11-12 mice per group for A-B; n=4 pools of 1-6 mice per group for C-G). Significance was tested by One-Way ANOVA with Tukey correction for multiple testing.

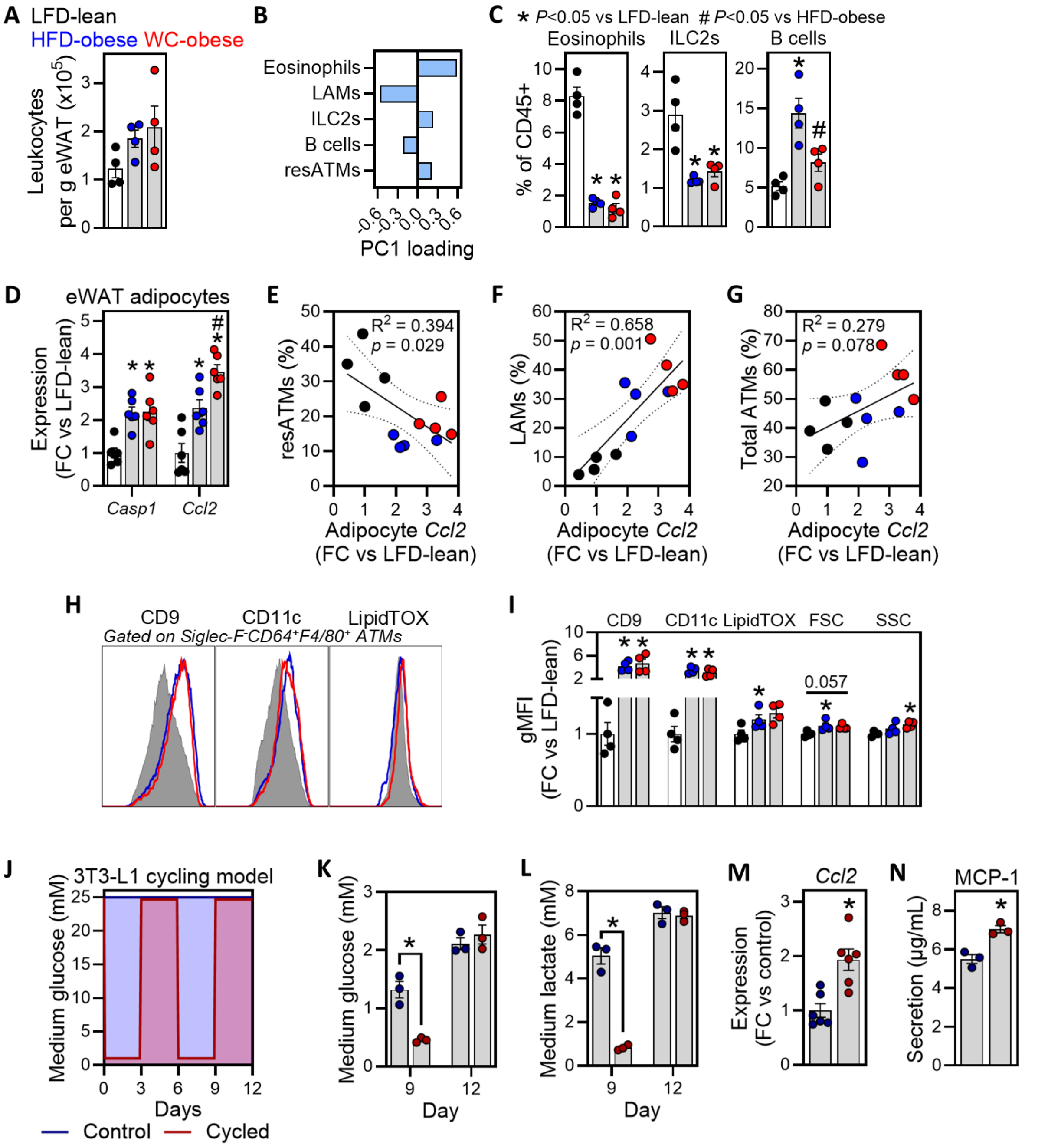


**Figure S6: Weight cycling increased gene expression and production of MCP-1 by adipocytes, which correlated with adipose tissue macrophages. Related to figure 6.**

(**A**) LFD-lean (black), HFD-obese (blue) and weight-cycled-obese (WC-obese, red) mice were sacrificed after a 10-week re-exposure of WC mice to HFD. Immune cells were isolated from epidydimal white adipose tissue (eWAT) and numbers per g tissue were determined. (**B**) Top 5 eWAT immune cell subsets contributing to the PC1 score of the PCA on the eWAT immune cell composition. (**C**) Frequencies of indicated eWAT immune cell subsets determined via flow cytometry. (**D**) Adipocytes were separated from the stromal vascular fraction of eWAT and adipocyte gene expression of the indicated markers was determined. (**E-G**) Correlation of adipocyte *Ccl2* expression with resident adipose tissue macrophages (resATMs; E), lipid-associated ATMs (LAMs; F) and total ATMs (G). (**H**) Model for nutrient cycling in 3T3-L1 adipocytes. (**I-J**) Medium glucose (I) and lactate (J) of 3T3-L1 adipocytes culture at indicated days. (**K-L**) Gene expression (K) and secretion (L) of MCP-1 by 3T3-L1 adipocytes at day 12. (**M-N**) Representative histograms of CD9,

CD11c and LipidTOX staining in adipose tissue macrophages (M) and quantification of indicated markers (N). Data are expressed as mean ± SEM or 95%CI (n=4-6 pools of 1-6 mice per group for A-G, M and N; n=3-6 technical replicates per group for H-L). Significance was tested by One-Way ANOVA with Tukey correction for multiple testing (A, C, D and N), F-test for the regression model (E-G) or paired t-test (I-L). * $p < 0.05$ vs LFD-lean, control, or as indicated, # $p < 0.05$ vs HFD-obese.

### Supplementary information

Supplementary Table 1: qPCR primers

| Gene | Forward primer | Reverse primer |
|---|---|---|
| *Acaca* | GATGAACCATCTCCGTTGGC | GACCCAATTATGAATCGGGAGTG |
| *Acox1* | TCGAAGCCAGCGTTACGAG | ATCTCCGTCTGGGCGTAGG |
| *Adgre1* | CTTTGGCTATGGGCTTCCAGTC | GCAAGGAGGACAGAGTTTATCGTG |
| *Adipoq* | GCAGAGATGGCACTCCTGGA | CCCTTCAGCTCCTGTCATTCC |
| *Casp1* | AATACAACCACTCGTACACGTC | AGCTCCAACCCTCGGAGAAA |
| *Ccl2* | CCCAATGAGTAGGCTGGAGA | TCTGGACCCATTCCTTCTTG |
| *Cd36* | AGATGACGTGGCAAAGAACAG | CCTTGGCTAGATAACGAACTCTG |
| *Cd68* | CCAATTCAGGGTGGAAGAAA | CTCGGGCTCTGATGTAGGTC |
| *Cidea* | TGACATTCATGGGATTGCAGAC | GGCCAGTTGTGATGACTAAGAC |
| *Clec4f* | AAAAGGCAAACGCTATGACTTCC | CCTCAACGCCTGGATCTCAG |
| *Col1a1* | TGTGTGCGATGACGTGCAAT | GGGTCCCTCGACTCCTACA |
| *Cpt1a* | CTCAGTGGGAGCGACTCTTCA | GGCCTCTGTGGTACACGACAA |
| *Fabp1* | GCCACCATGAACTTCTCCGGCA | GGTCCTCGGGCAGACCTATTGC |
| *Il1b* | GACCCCAAAAGATGAAGGGCT | ATGTGCTGCTGCGAGATTTG |
| *Il1ra* | AAATCTGCTGGGGACCCTAC | TGAGCTGGTTGTTTCTCAGG |
| *Il6* | AGTCCTTCCTACCCCAATTTCC | TTGGTCCTTAGCCACTCCTTC |
| *Insr* | CGAGTGCCCGTCTGGCTATA | GGCAGGGTCCCAGACATG |
| *Itgax* | CTGGATAGCCTTTCTTCTGCTG | GCACACTGTGTCCGAACTCA |
| *Lcn2* | TGGAAGAACCAAGGAGCTGT | GGTGGGGACAGAGAAGA |
| *Lep* | AGAAGATCCCAGGGAGGAAA | TGATGAGGGTTTTGGTGTCA |
| *Pnpla2* | CAACGCCACTCACATCTACGG | GGACACCTCAATAATGTTGGCAC |
| *36b4* | ATGGGTACAAGCGCGTCCTG | GCCTTGACCTTTTCAGTAAG |
| *Scd1* | TAGCCTGTAAAAGATTTCTGCAAACC | CCGGAGACCCTTAGATCGA |
| *Srebp1* | GGAGCCATGGATTGCACATT | CCTGTCTCACCCCCAGCATA |
| *Timp1* | GCAACTCGGACCTGGTCATAA | CGGCCCGTGATGAGAAACT |
| *Tnf* | CCCTCACACTCAGATCATCTTCT | GCTACGACGTGGGCTACAG |
| *Trem2* | CTGGAACCGTCACCATCACTC | CGAAACTCGATGACTCCTCGG |

Supplementary Table 2: Antibodies and reagents for flow cytometry

| **Target** | **Clone** | **Conjugate** | **Source** | **Identifier** |
|---|---|---|---|---|
| B220 | RA3-6B2 | Alexa Fluor 700 | Biolegend | 103231 |
| B220 | RA3-6B2 | BV510 | Biolegend | 103247 |
| B220 | RA3-6B2 | BV650 | Biolegend | 103241 |
| CD3 | 17A2 | BV650 | Biolegend | 100229 |
| CD3 | 17A2 | FITC | Biolegend | 100203 |
| CD4 | RM4-5 | BV785 | Biolegend | 100551 |
| CD8 | 53-6.7 | PE-Cy7 | Biolegend | 100721 |
| CD8 | 53-6.7 | PerCP-Cy5.5 | Biolegend | 100733 |
| CD9 | MZ3 | APC | Biolegend | 124811 |
| CD11b | M1/70 | APC | Biolegend | 101211 |
| CD11b | M1/70 | PerCP-Cy5.5 | Biolegend | 101227 |
| CD11c | N418 | BV605 | Biolegend | 117333 |
| CD16/32 | 93 | - | Biolegend | 101319 |
| CD25 | PC61 | PE-Dazzle 594 | Biolegend | 102047 |
| CD44 | IM7 | BV605 | Biolegend | 103047 |
| CD45 | 30-F11 | APC-Fire 750 | Biolegend | 103153 |
| CD45 | 30-F11 | Pacific Blue | Biolegend | 103125 |
| CD62L | MEL-14 | PE-Cy7 | Biolegend | 104417 |
| CD64 | X54-5/7.1 | PE-Cy7 | Biolegend | 139313 |
| CLEC2 | 17D9 | FITC | Bio-RAD | MCA5700 |
| F4/80 | BM8 | PE | Biolegend | 123109 |
| FOXP3 | MF-14 | Alexa Fluor 647 | Biolegend | 126407 |
| Ly6C | HK1.4 | Alexa Fluor 700 | Biolegend | 128023 |
| Ly6G | 1A8 | PerCP-Cy5.5 | Biolegend | 127615 |
| MHCII | M5/114.15.2 | BV785 | Biolegend | 107645 |
| NK1.1 | PK136 | BV650 | Biolegend | 108735 |
| Siglec-F | S17007L | PE-Dazzle 594 | Biolegend | 155529 |
| TCRγδ | UC7-13D5 | PE | Biolegend | 107507 |
| TIM4 | RMT4-54 | PE | Biolegend | 130005 |
| **Reagent** | | | **Source** | **Identifier** |
| HCS LipidTOX™ Green Neutral Lipid Stain | | | Thermo Fisher | H34475 |
| Zombie Aqua™ Fixable Viability Kit | | | Biolegend | 423101 |
